# Transcriptomic data of larval zebrafish exposed to continuous sub- and supra-MCL sodium arsenite and uranyl nitrate

**DOI:** 10.64898/2026.02.22.707205

**Authors:** Phillip H. Kalaniopio, Ronald S. Allen, Matthew C. Salanga

## Abstract

Uranium (U) and arsenic (As) are both ubiquitous contaminants in the American southwest, posing risks to humans, animals, and the environment. Depleted uranium’s (DU) chronic effects and mechanisms of toxicity are incompletely understood. Differential gene expression of concomitant exposures to identify markers of toxicity have not been undertaken until now. Continuous low-dose, high-dose, and concomitant exposures are investigated using the larval zebrafish (*Danio rerio*), with exposure paradigms lasting from embryo collection until sampling at 5 days post fertilization (dpf). Herein, we describe overall differential gene expression with counts and pathway enrichment statistics using both gene ontology (GO) and Kyoto Encyclopedia of Genes and Genomes (KEGG) analyses. The raw dataset has been deposited in NCBI’s Gene Expression Omnibus (GEO) repository [1] under the accession number *GSE319292* [2].

| Specification Table |  |
| --- | --- |
| Subject area | Omics - Transcriptomics |
| More specific subject area | Bioinformatics and toxicogenomics of metal contaminants at environmentally relevant levels |
| Type of data | Figures, tables – processed and analyzed |
| How data was acquired | High-throughput RNA sequencing |
| Data format | Filtered and analyzed with statistical tests |
| Experimental factors | Wildtype AB larval zebrafish exposed to sodium arsenite and uranyl nitrate at environmentally relevant levels |
| Experimental features | Comparison of the transcriptomic responses from pooled larval zebrafish exposed to either 1 or 10 μM sodium arsenite, 30 or 300 ppb uranyl nitrate, or 1 μM sodium arsenite plus 30 ppb uranyl nitrate in combination, to a vehicle media control. Biological quadruplicates were incorporated for each condition. |
| Data source location | Department of Biological Sciences, Northern Arizona University, 86011, AZ, USA |
| Data accessibility | Data are available via NCBI's Gene Expression Omnibus (GEO) accession number GSE319292<br>https://www.ncbi.nlm.nih.gov/geo/query/acc.cgi?acc=GSE319292 |
| Related research article | None |

**VALUE OF THE DATA:**

- Uranyl nitrate (UN), a water-soluble depleted uranium species, and sodium arsenite (As) are both ubiquitous contaminants in the American southwest, posing risks to humans, animals, and the environment. The United States Environmental Protection Agency (EPA) has set maximum contaminant limits (MCL) of 30 ppb U atoms and 10 ppb As atoms, respectively.
- These data show differentially expressed genes (DEGs) from larval zebrafish exposed to 1 or 10 µM As, 30 or 300 µg/L UN, or 1 µM As and 30 µg/L UN in combination. Concentrations were specifically chosen based on environmental relevance.
- Gene ontology (GO) and Kyoto Encyclopedia of Genes and Genomes (KEGG) pathway enrichment analyses of up- and down-regulated DEGs are provided to understand the molecular mechanisms of uranium toxicity and inform future studies.
- These data should be used for biomarker identification and mechanistic interrogation of single and combinatorial exposures of environmentally relevant compounds at realistic exposure levels.

## BACKGROUND

Uranium and arsenic are environmental contaminants of concern in the American southwest. The EPA set the MCLs for drinking water at 30 parts per billion (ppb; 30 µg/L; 0.126 µM) of U atoms and 10 ppb As (10 µg/L; 0.077 µM) [3]. Credo et al. collected samples from wells across the Navajo Nation and quantified contaminants using ICP-MS. For U, 21 of 231 wells exceeded the MCL, 10 were near the MCL, and 179 were below the MCL but still had detectable U. For As, 40 of 235 wells exceeded the 10 ppb MCL, while 157 were below the MCL yet above the detection limit [4]. There are limited *in vivo* toxicological studies at levels below the MCL for both metals, and there is even less transcriptomic data of samples exposed to environmentally relevant levels of these contaminants, both above and below the MCL. Armant and company performed transcriptomic studies of DU-exposure in zebrafish and samples consisted of ovary, testis, and brain tissue from adults following a 10 day exposure regime and the 2-cell and 96 hour post-fertilization (hpf) progeny of those exposed fish (data accessible at NCBI GEO database, accession *GSE96603*) [5], but were limited to 20 µg/L UN exposure doses. Here, we seek to provide a transcriptomic resource for *in vivo* sub- and supra-MCL exposures of U and As individually and concomitantly at sub-MCL exposure, using the larval zebrafish (*Danio rerio*).

## DATA DESCRIPTION

These data contain 24 paired-end high-throughput RNA sequencing samples of 5 dpf larvae exposed to either: 1 or 10 µM sodium arsenite (sample names AsL and AsH, respectively), 30 or 300 µg/L uranyl nitrate (UL or UH, respectively), or 1 µM sodium arsenite and 30 µg/L uranyl nitrate in combination (AsU). The full list of these samples, including genotype and background, number of larvae per sample, and exposure compounds and concentrations, is provided in **Table 1**, showing an average of 58,907,817.2 clean reads per sample, resulting in 8.83 Gb of clean base calls, and an average Q30 Phred score of 94.65% (**Table 1**). Counts of differentially expressed genes are described per condition versus vehicle control using volcano scatterplots and are tabulated (**Figures 1a-e**; **Table 2**). Overlapping expression is displayed in the Venn diagram presenting the number of genes that are uniquely expressed within each sample, with the overlapping regions showing the number of genes that are expressed in two or more samples (**Figure 1f**). The top five up- and down-regulated differentially expressed genes per treatment condition versus vehicle control are provided in (**Table 3**). The top five up- and down-regulated gene ontology (GO) terms with term type and associated p-values per treatment condition versus vehicle control are provided in (**Table 4**). The top five up- and down-regulated KEGG terms with raw p-values, corrected p-values, and contributing KEGG IDs per treatment condition versus vehicle control are provided in (**Table 5**). Scatterplots showing KEGG pathway enrichment results compared to control are drawn in (**Figure 2a-e**).

**Table 1.**
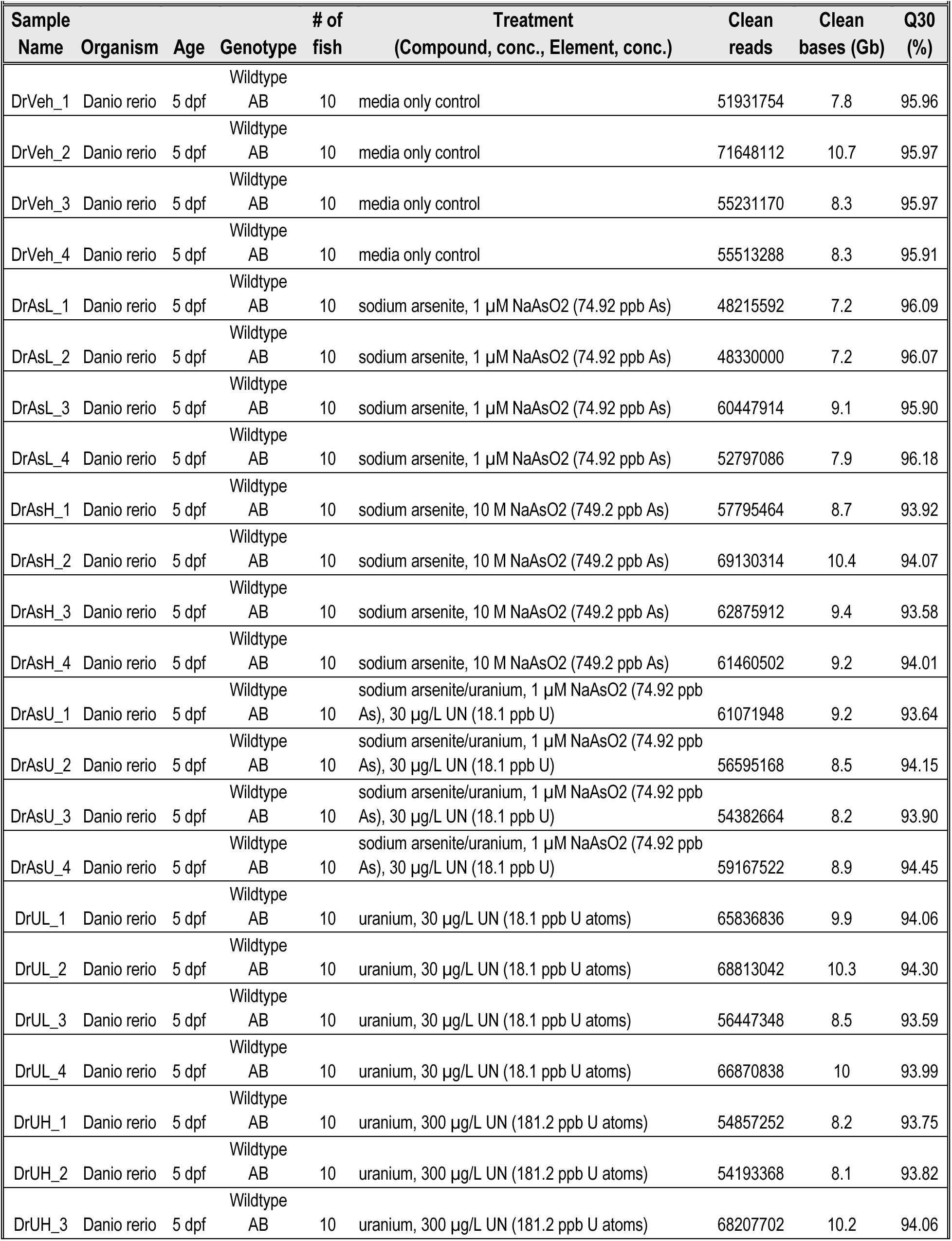
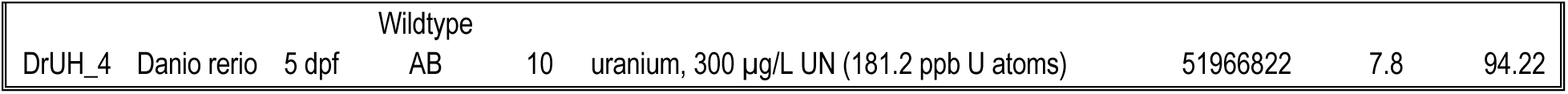
Sample information of HTS RNA-sequencing experiment. Columns include sample name for ease of reference, organism, age, genotype, number of fish pooled in sample, the treatment compounds, concentrations, and elements, and resulting clean reads, bases in gigabase (Gb) and Q30 Phred scores as percentages.

**Figure 1a.**
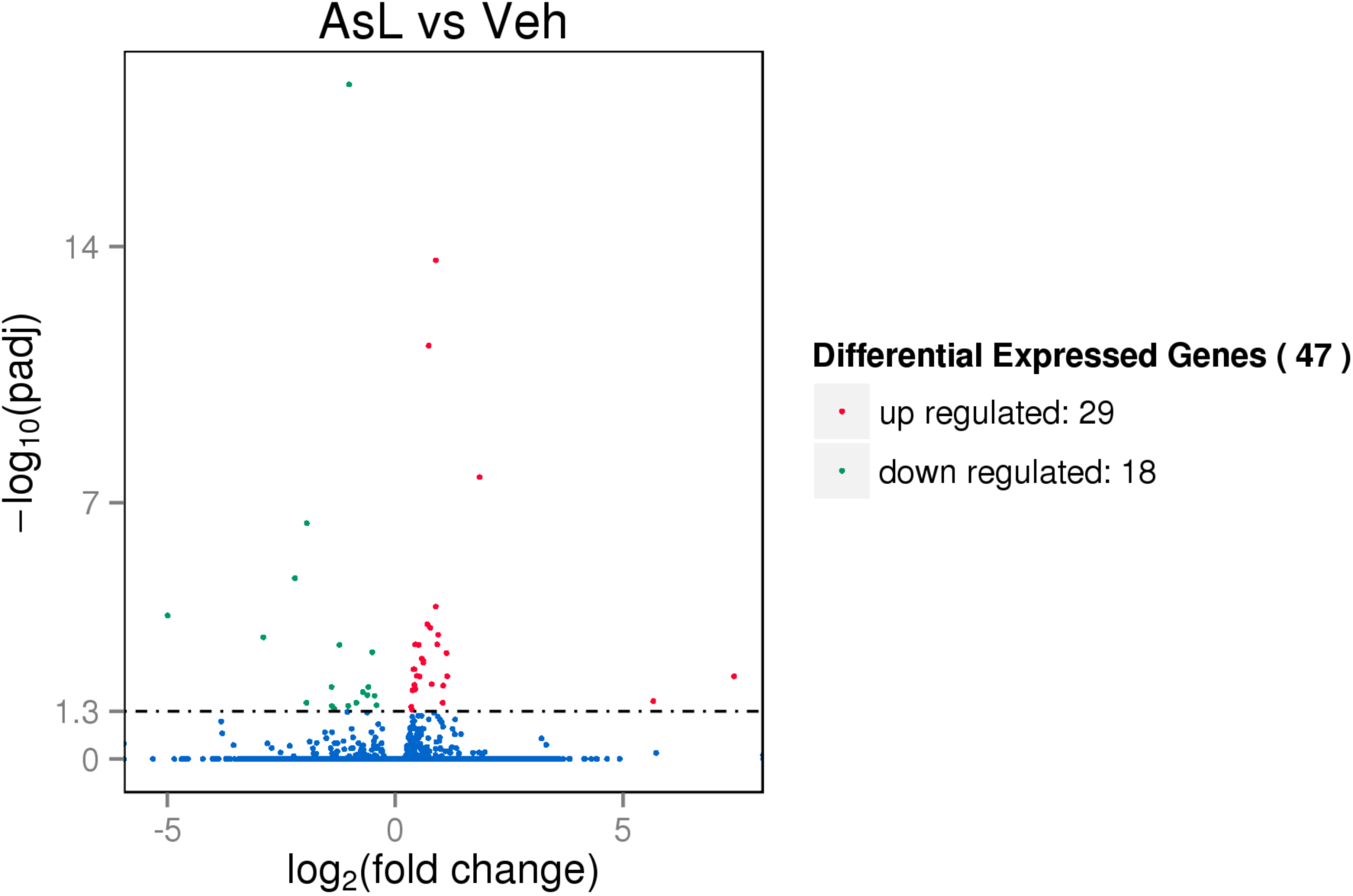
AsL vs. Veh DEG volcano plot.

**Figure 1b.**
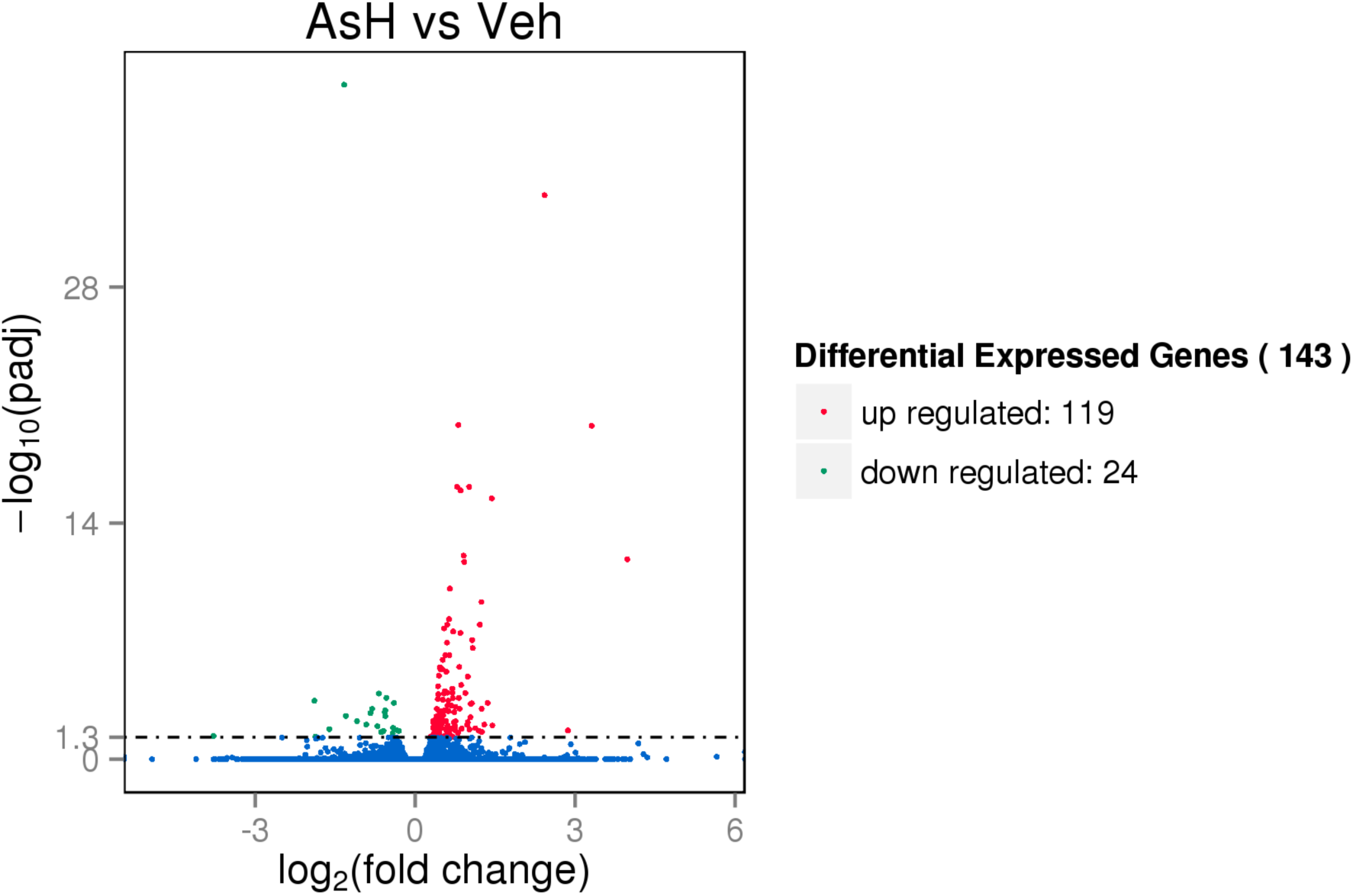
AsH vs. Veh DEG volcano plot.

**Figure 1c.**
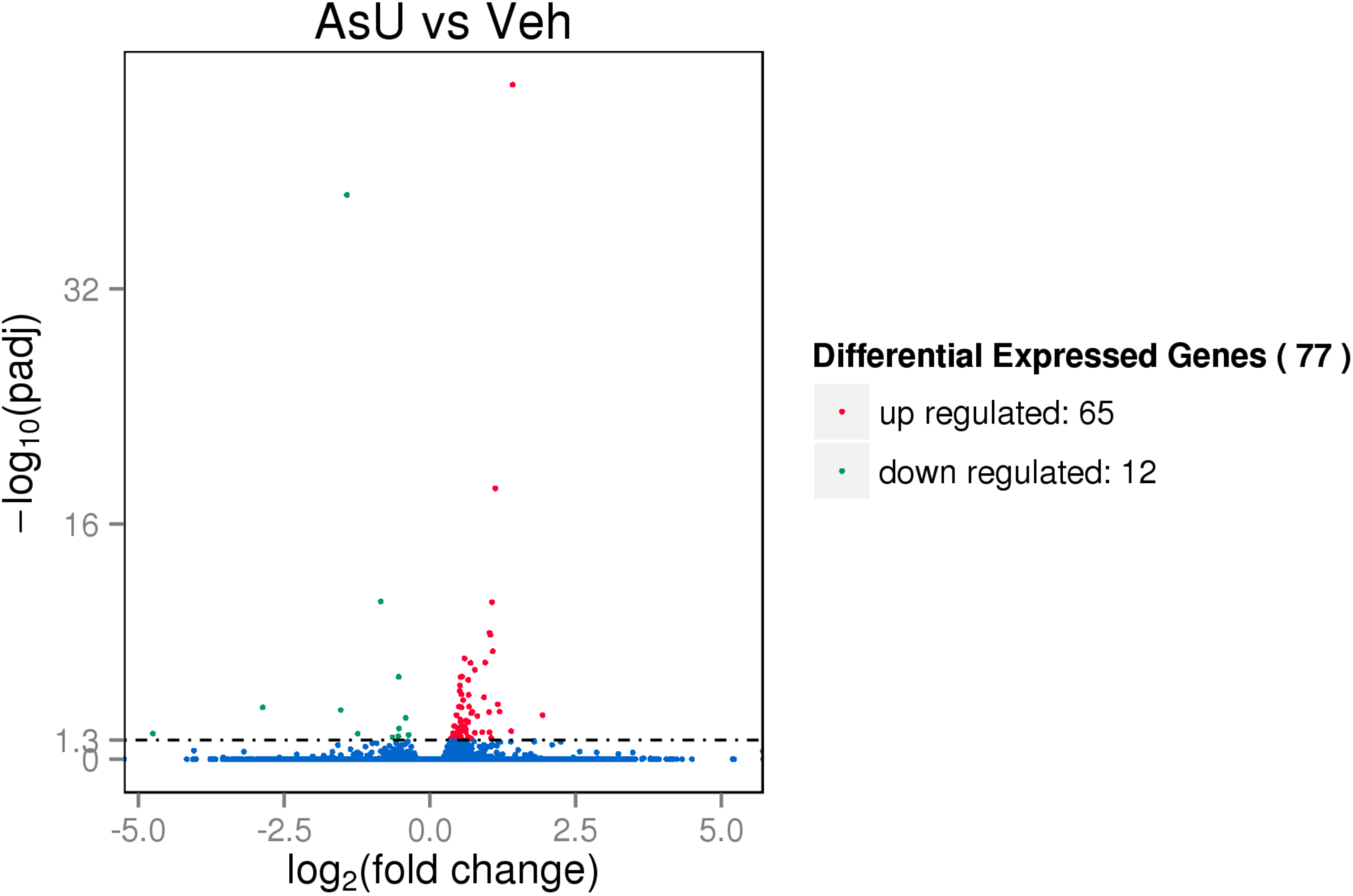
AsU vs. Veh DEG volcano plot.

**Figure 1d.**
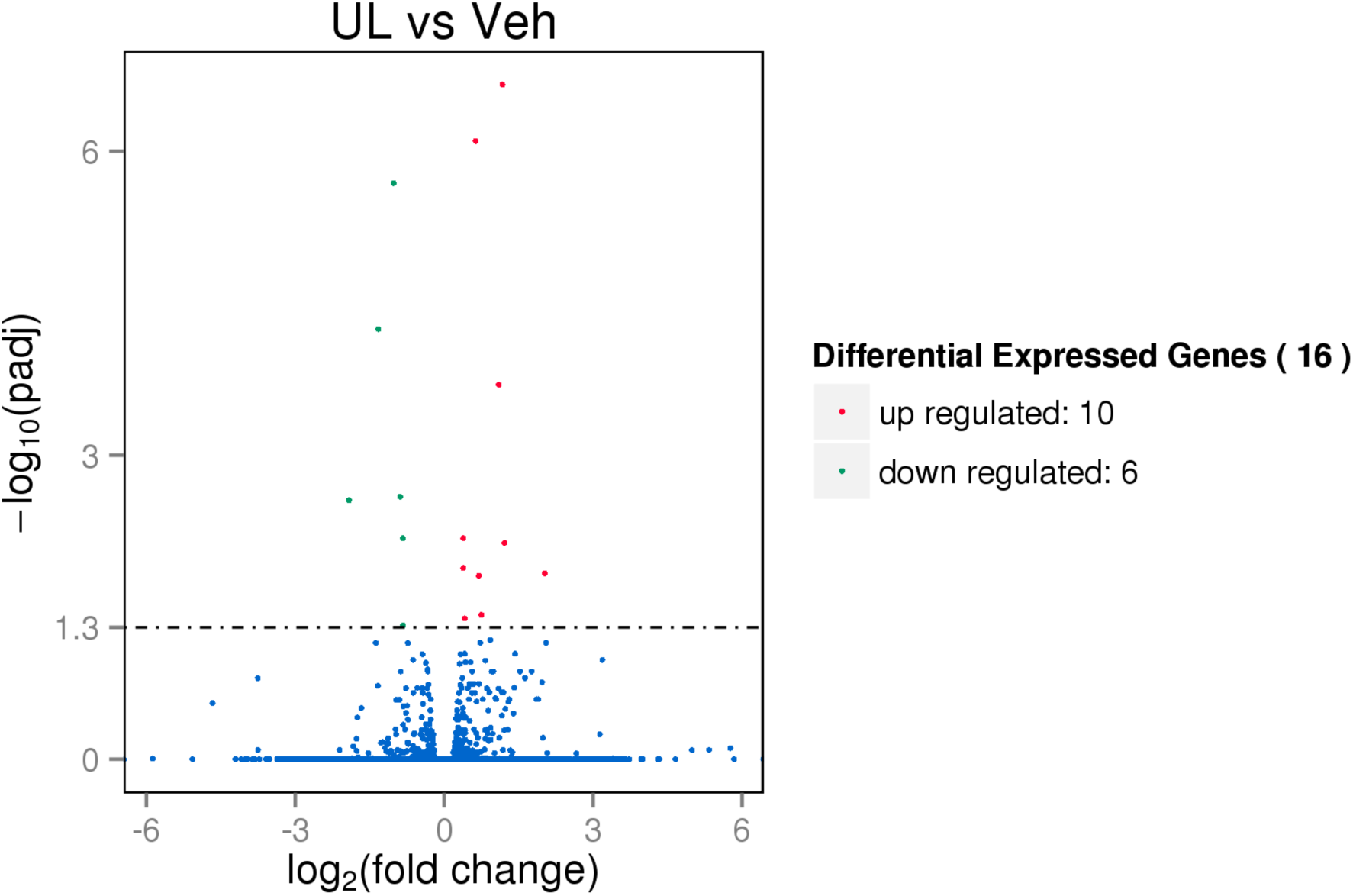
UL vs. Veh DEG volcano plot.

**Figure 1e.**
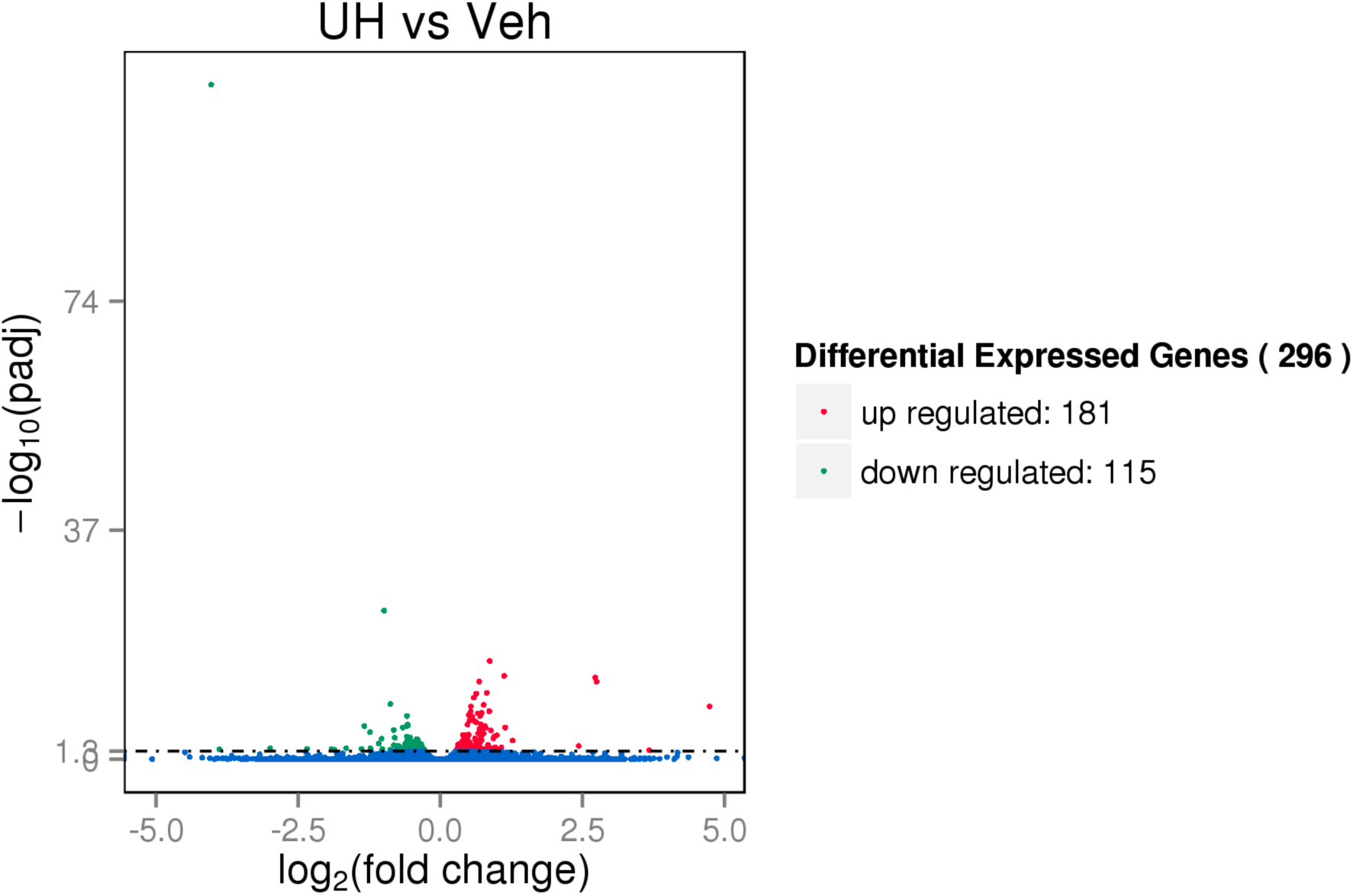
UH vs. Veh DEG volcano plot.

**Figure 1f.**
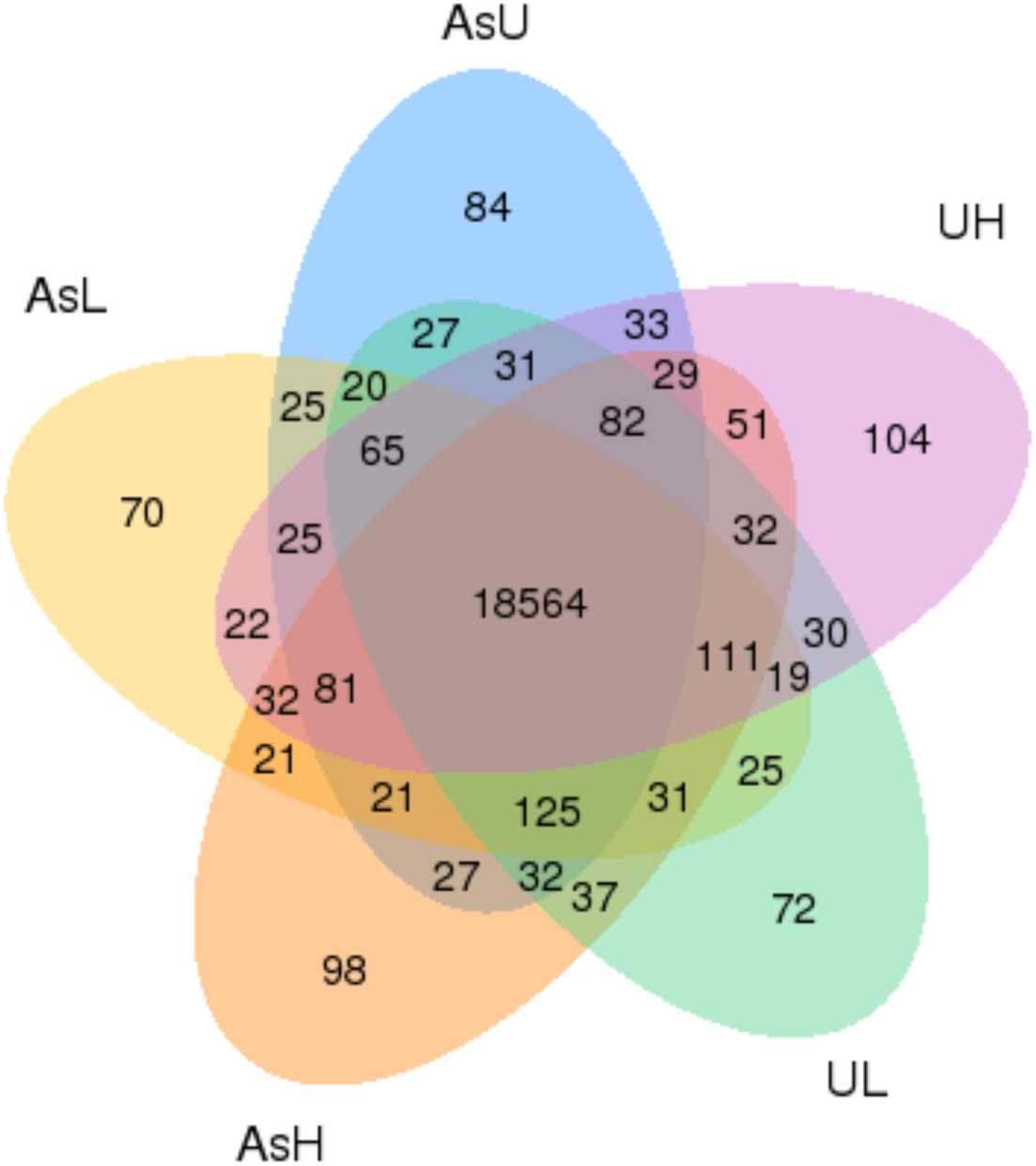
Co-expression Venn diagram of DEGs in treatment groups.

**Table 2.** Number of differentially expressed genes per treatment group compared to vehicle control (p_adj_ > 0.05).

| **Table 2.** Overall differential gene expression. |  |  |  |
| --- | --- | --- | --- |
| **Treatment**<br>**(vs. Vehicle)** | **Up-regulated** | **Down-regulated** | **Total** |
| AsL | 29 | 18 | 47 |
| AsH | 119 | 24 | 143 |
| AsU | 65 | 12 | 77 |
| UL | 10 | 6 | 16 |
| UH | 181 | 115 | 296 |

**Table 3.** Top five differentially expressed genes per treatment group, separated by up-or down-regulation, versus vehicle control with BLAST name, Gene ID, log_2_ fold change, p-value, and adjusted p-value.

| Table 3. Top five DEGs per treatment group compared to vehicle control. |  |  |  |  |  |
| --- | --- | --- | --- | --- | --- |
| Gene set (vs. vehicle) | Blast Name | Gene ID | Log<sub>2</sub> Fold Change | p-value | p-adjusted |
| AsL Up | MIA | 569807 | 7.4358 | 4.84E-06 | 0.0055002 |
|  | DNJ6A | 393275 | 5.6635 | 3.40E-05 | 0.026459 |
|  | PSME3 | 30649 | 1.853 | 2.71E-12 | 2.00E-08 |
|  | SP2 | 556835 | 1.1397 | 4.75E-06 | 0.0055002 |
|  | KCNJ1 | 569052 | 1.1244 | 7.80E-07 | 0.0012803 |
| AsL Down | LOC108190702 | 108190702 | -4.9924 | 3.28E-08 | 0.00012106 |
|  | SH3R3 | 100537944 | -2.8929 | 1.93E-07 | 0.00047541 |
|  | ADH1 | 402841 | -2.2011 | 2.35E-09 | 1.16E-05 |
|  | PTPA | 445031 | -1.9456 | 4.00E-05 | 0.029237 |
|  | DAPK3 | 571352 | -1.9412 | 6.15E-11 | 3.63E-07 |
| AsH Up | MANF | 767691 | 3.9779 | 4.70E-16 | 1.40E-12 |
|  | YCP6 | 641422 | 3.3092 | 2.25E-24 | 1.67E-20 |
|  | TRI39 | 110437745 | 2.8652 | 8.27E-05 | 0.019806 |
|  | PTGR1 | 566996 | 2.4266 | 2.41E-38 | 3.57E-34 |
|  | PSME3 | 30649 | 1.4481 | 3.54E-05 | 0.010117 |
| AsH Down | PBDC1 | 436977 | -3.7836 | 0.00019879 | 0.042485 |
|  | LADD | 100002541 | -1.8905 | 5.85E-07 | 0.00034104 |
|  | SH3R3 | 100537944 | -1.8689 | 0.00021735 | 0.045793 |
|  | CSN8 | 393198 | -1.6087 | 6.79E-05 | 0.016878 |
|  | SERPH | 30449 | -1.3318 | 3.36E-45 | 9.99E-41 |
| AsU Up | AMER1 | 100885784 | 1.9341 | 1.16E-06 | 0.0010038 |
|  | ARRD3 | 447866 | 1.4207 | 4.42E-51 | 1.30E-46 |
|  | ATNG | 794522 | 1.3925 | 2.06E-05 | 0.01238 |
|  | PRL | 791526 | 1.1975 | 5.94E-07 | 0.0005847 |
|  | SMS2 | 79186 | 1.1648 | 1.45E-07 | 0.00018617 |
| AsU Down | RPAC2 | 445412 | -4.7518 | 3.53E-05 | 0.018292 |
|  | SH3R3 | 100537944 | -2.8659 | 2.78E-07 | 0.00030412 |
|  | SRAC1 | 568022 | -1.5304 | 4.33E-07 | 0.00045593 |
|  | SERPH | 30449 | -1.422 | 2.78E-43 | 4.10E-39 |
|  | FBX4 | 100319387 | -1.2369 | 3.67E-05 | 0.018673 |
| UL Up | AMER1 | 100885784 | 2.0237 | 5.94E-06 | 0.014645 |
|  | CSL3 | 449685 | 1.2145 | 2.48E-06 | 0.0073333 |
|  | YSN9 | 100034462 | 1.1734 | 7.45E-12 | 2.21E-07 |
| Gene set (vs. vehicle) | Blast Name | Gene ID | Log<sub>2</sub> Fold Change | p-value | p-adjusted |
|  | VGFR1 | 544667 | 1.1003 | 3.41E-08 | 0.00020194 |
|  | FABPI | 30708 | 0.74934 | 1.78E-05 | 0.037731 |
| UL Down | LADD | 100002541 | -1.9192 | 6.59E-07 | 0.0027885 |
|  | LOC110438463 | 110438463 | -1.3271 | 7.69E-09 | 5.69E-05 |
|  | MFSD8 | 564342 | -1.0216 | 2.09E-10 | 2.06E-06 |
|  | CP4V2 | 562008 | -0.88477 | 5.21E-07 | 0.0025699 |
|  | LOC108190590 | 108190590 | -0.83194 | 1.98E-06 | 0.006592 |
| UH Up | SSH | 101886694 | 4.7362 | 1.42E-12 | 3.20E-09 |
|  | ADCK2 | 572087 | 3.676 | 0.00032882 | 0.037292 |
|  | CC115 | 503608 | 2.7516 | 7.04E-17 | 2.98E-13 |
|  | TRI39 | 110437745 | 2.7268 | 1.11E-17 | 6.58E-14 |
|  | SCF | 553181 | 2.435 | 2.98E-05 | 0.0073044 |
| UH Down | CSN8 | 393198 | -4.0312 | 3.57E-114 | 1.06E-109 |
|  | PBDC1 | 436977 | -3.8835 | 0.00017517 | 0.025171 |
|  | MX1 | 561067 | -2.9908 | 0.00010625 | 0.017769 |
|  | TLS1 | 449772 | -2.3444 | 0.0001372 | 0.021278 |
|  | CR318588.1 | 100535167 | -1.9097 | 0.00016833 | 0.024546 |

**Table 4.** Top five enriched gene ontology (GO) terms, both up- and down-regulated, per treatment condition compared to vehicle control including term type (biological_process, molecular_function, or cellular_component), p-value, and corrected p-value.

| **Table 4.** Top five up- and down-regulated GO terms per treatment group compared to vehicle control. |  |  |  |  |
| --- | --- | --- | --- | --- |
| **Gene set (vs. vehicle)** | **GO Term** | **Type** | **p-value** | **Corrected p-value** |
| **AsL Up** | response to lipid | biological_process | 0.0022099 | 1 |
|  | steroid hormone mediated signaling pathway | biological_process | 0.0022099 | 1 |
|  | response to steroid hormone | biological_process | 0.0022099 | 1 |
|  | cellular response to steroid hormone stimulus | biological_process | 0.0022099 | 1 |
|  | cellular response to lipid | biological_process | 0.0022099 | 1 |
| **AsL Down** | phosphatase activator activity | molecular_function | 0.0011271 | 1 |
|  | CDP-diacylglycerol diphosphatase activity | molecular_function | 0.001332 | 1 |
|  | protein import into nucleus | biological_process | 0.001477 | 1 |
|  | protein localization to nucleus | biological_process | 0.001477 | 1 |
|  | protein targeting to nucleus | biological_process | 0.001477 | 1 |
| **AsH Up** | oxidoreductase activity | molecular_function | 8.24E-08 | 0.00039488 |
|  | cofactor binding | molecular_function | 5.16E-06 | 0.010206 |
|  | coenzyme binding | molecular_function | 6.39E-06 | 0.010206 |
|  | selenium binding | molecular_function | 1.77E-05 | 0.019784 |
|  | oxidation-reduction process | biological_process | 2.06E-05 | 0.019784 |
| **AsH Down** | extracellular space | cellular_component | 6.20E-05 | 0.17609 |
|  | fibrinogen complex | cellular_component | 0.000238 | 0.17609 |
|  | platelet activation | biological_process | 0.000238 | 0.17609 |
|  | blood coagulation | biological_process | 0.00028053 | 0.17609 |
|  | cell activation | biological_process | 0.00033321 | 0.17609 |
| **AsU Up** | sequence-specific DNA binding | molecular_function | 2.95E-08 | 9.37E-05 |
|  | nucleic acid binding transcription factor activity | molecular_function | 5.86E-08 | 9.37E-05 |
|  | transcription factor activity, sequence-specific DNA binding | molecular_function | 5.86E-08 | 9.37E-05 |
|  | regulation of primary metabolic process | biological_process | 4.75E-05 | 0.031934 |
|  | regulation of transcription, DNA-templated | biological_process | 6.63E-05 | 0.031934 |
| **AsU Down** | detection of visible light | biological_process | 0.0020188 | 1 |
|  | protein-chromophore linkage | biological_process | 0.0023607 | 1 |
|  | detection of external stimulus | biological_process | 0.0027052 | 1 |
|  | detection of abiotic stimulus | biological_process | 0.0027052 | 1 |
|  | detection of light stimulus | biological_process | 0.0027052 | 1 |
| **UL Up** | voltage-gated calcium channel activity | molecular_function | 0.0027785 | 1 |
|  | voltage-gated calcium channel complex | cellular_component | 0.0040412 | 1 |
| **Gene set**<br>**(vs.**<br>**vehicle)** | **GO Term** | **Type** | **p-value** | **Corrected**<br>**p-value** |
|  | calcium channel complex | cellular_component | 0.0040412 | 1 |
|  | calcium ion transmembrane transport | biological_process | 0.00681 | 1 |
|  | ion channel complex | cellular_component | 0.010063 | 1 |
| **UL Down** | heme binding | molecular_function | 0.024837 | 1 |
|  | tetrapyrrole binding | molecular_function | 0.027031 | 1 |
|  | iron ion binding | molecular_function | 0.028928 | 1 |
|  | oxidoreductase activity, acting on paired donors, with incorporation or reduction of molecular oxygen | molecular_function | 0.029199 | 1 |
|  | DNA-directed RNA polymerase activity | molecular_function | 0.035992 | 1 |
| **UH Up** | sequence-specific DNA binding | molecular_function | 1.31E-06 | 0.0062775 |
|  | nucleic acid binding transcription factor activity | molecular_function | 7.84E-06 | 0.01113 |
|  | transcription factor activity, sequence-specific DNA binding | molecular_function | 7.84E-06 | 0.01113 |
|  | anion transmembrane transporter activity | molecular_function | 1.01E-05 | 0.01113 |
|  | organic anion transmembrane transporter activity | molecular_function | 1.16E-05 | 0.01113 |
| **UH Down** | hydrolase activity, acting on acid anhydrides, in phosphorus-containing anhydrides | molecular_function | 2.52E-09 | 6.72E-06 |
|  | hydrolase activity, acting on acid anhydrides | molecular_function | 2.81E-09 | 6.72E-06 |
|  | nucleoside-triphosphatase activity | molecular_function | 2.30E-08 | 3.59E-05 |
|  | pyrophosphatase activity | molecular_function | 3.00E-08 | 3.59E-05 |
|  | DNA replication | biological_process | 4.00E-08 | 3.83E-05 |

**Table 5.**
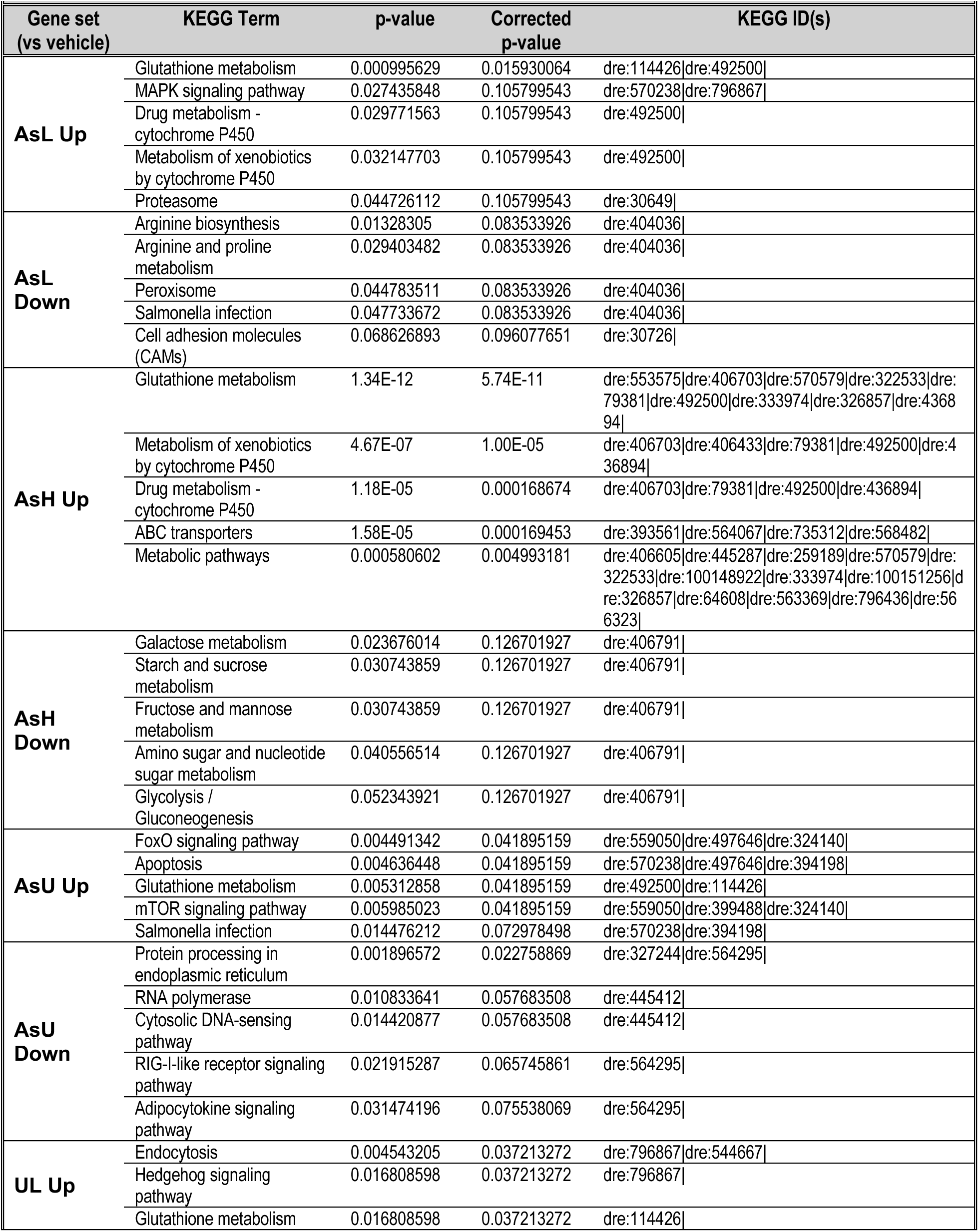
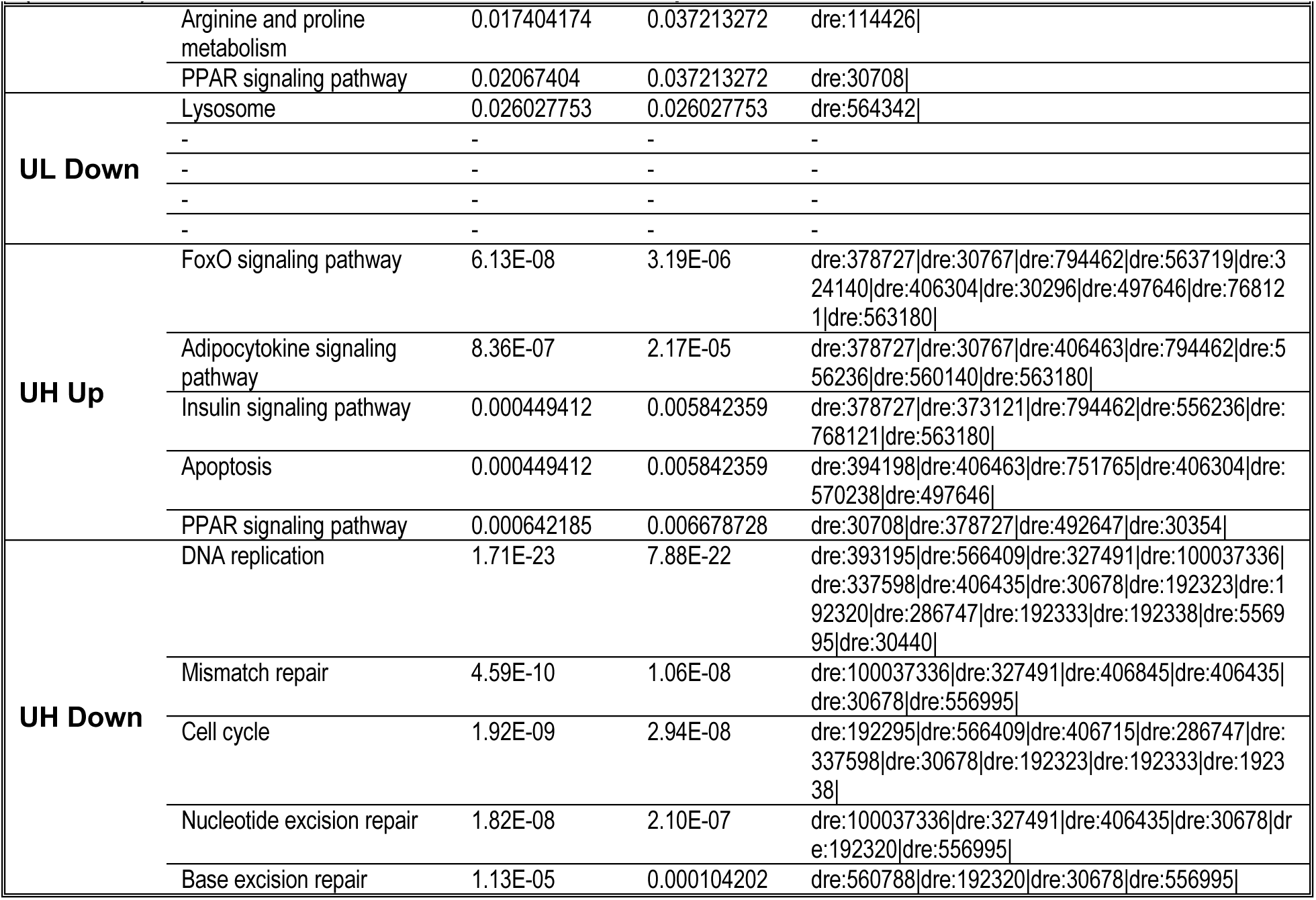
Top five enriched KEGG terms per treatment condition compared to vehicle control including term, p-value, corrected p-value, and contributory KEGG IDs.

**Figure 2a.**
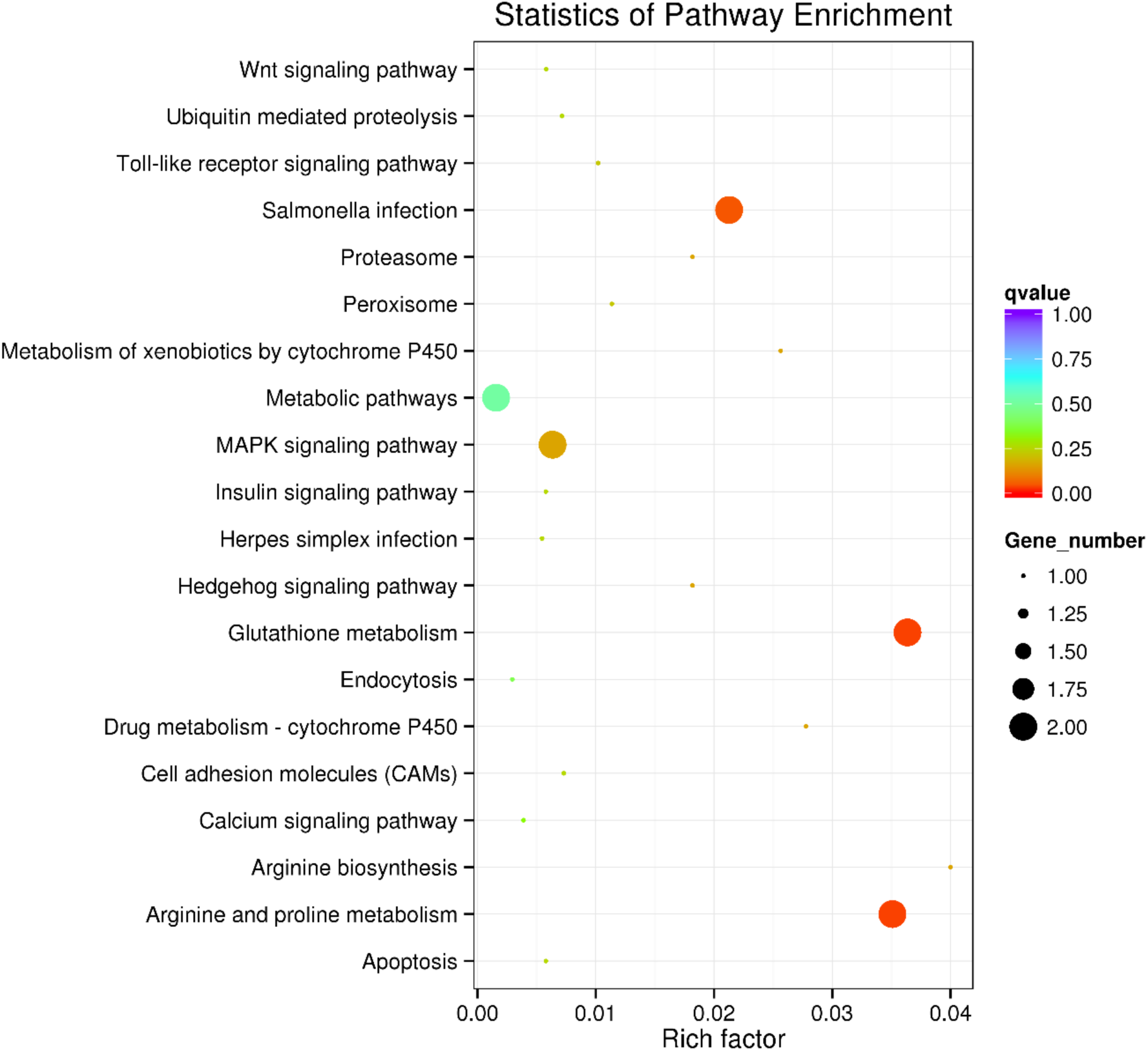
Overall KEGG pathway enrichment statistics for AsL vs Veh.

**Figure 2b.**
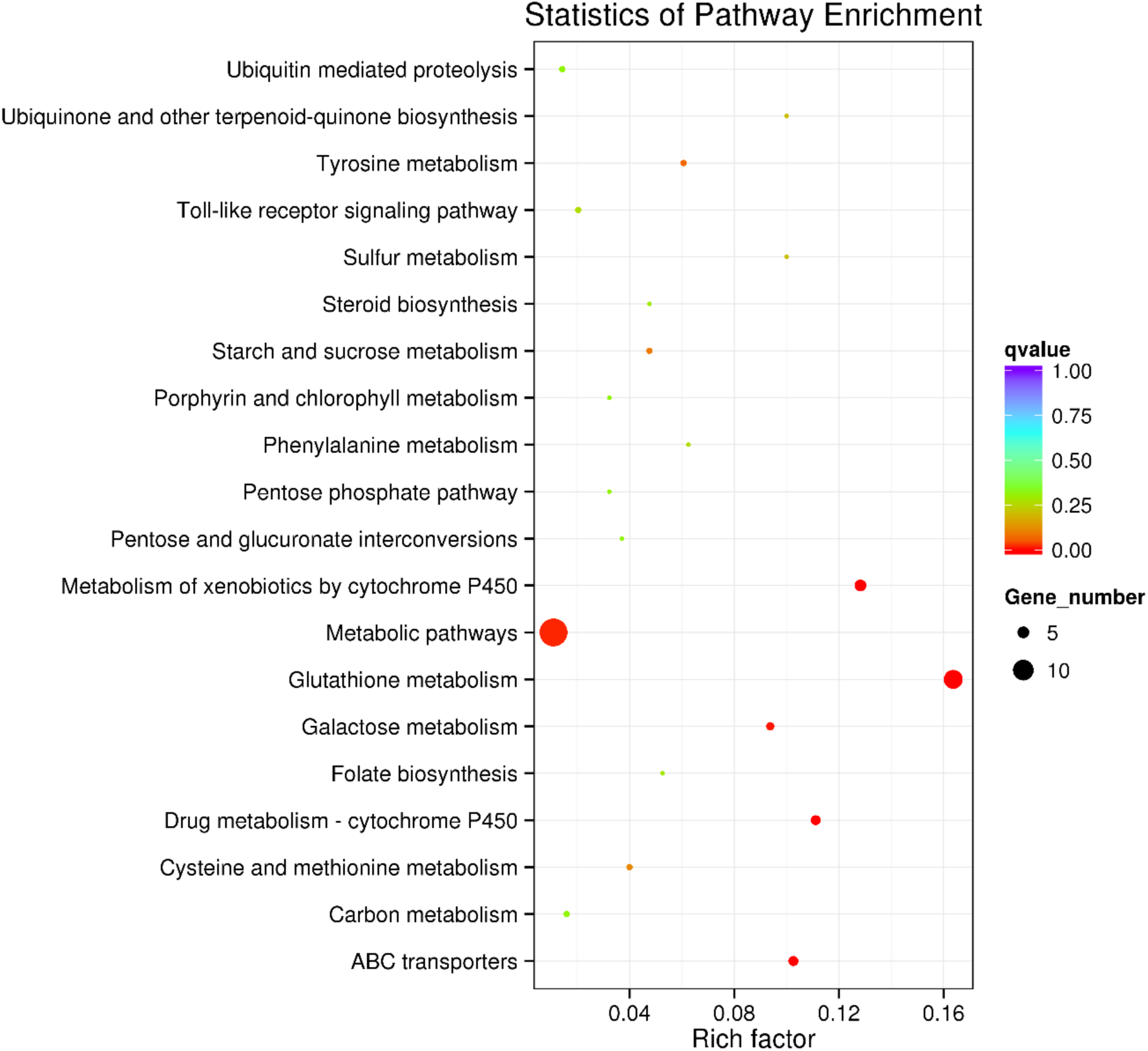
Overall KEGG pathway enrichment statistics for AsH vs Veh.

**Figure 2c.**
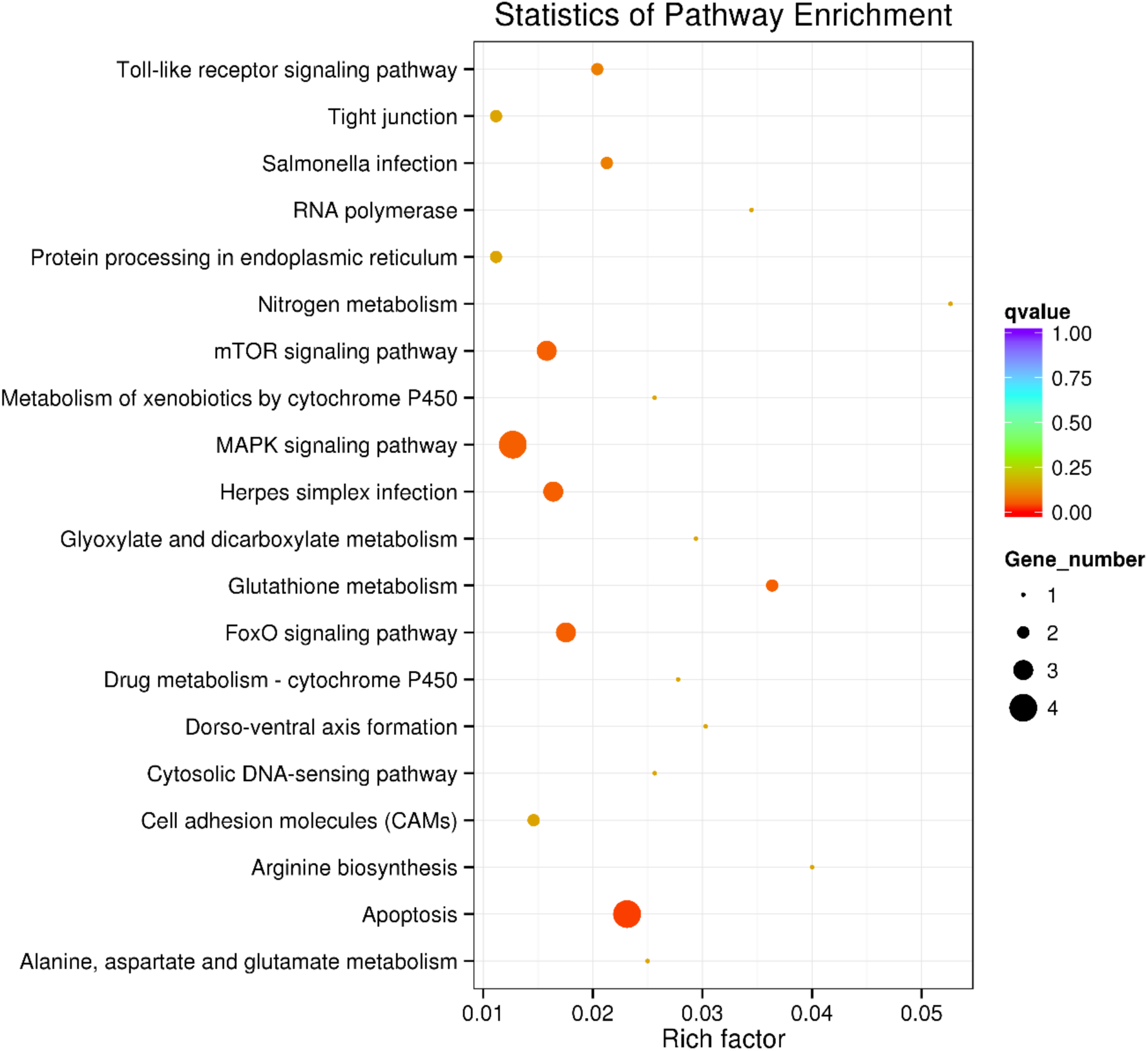
Overall KEGG pathway enrichment statistics for AsU vs Veh.

**Figure 2d.**
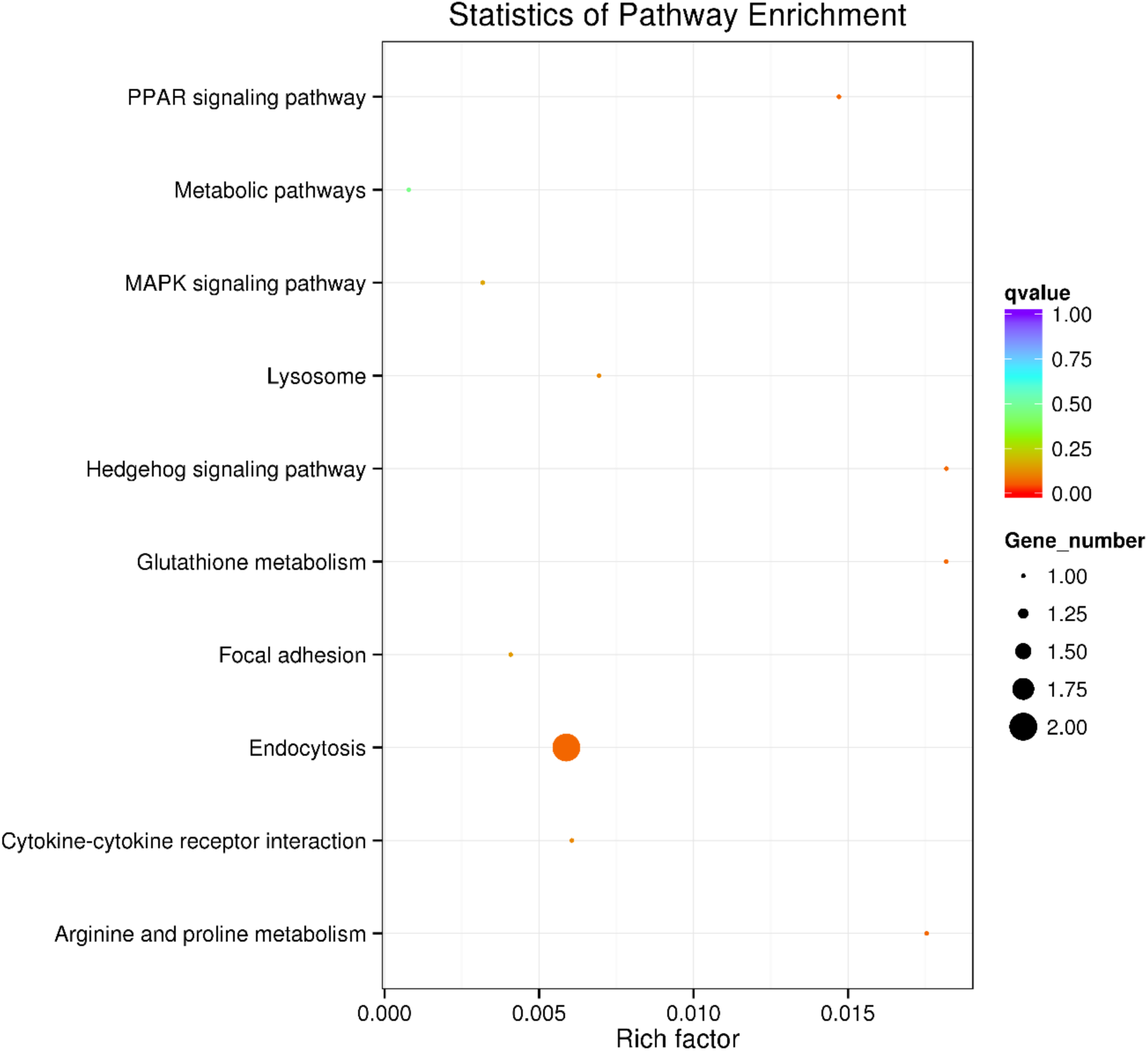
Overall KEGG pathway enrichment statistics for UL vs Veh.

**Figure 2e.**
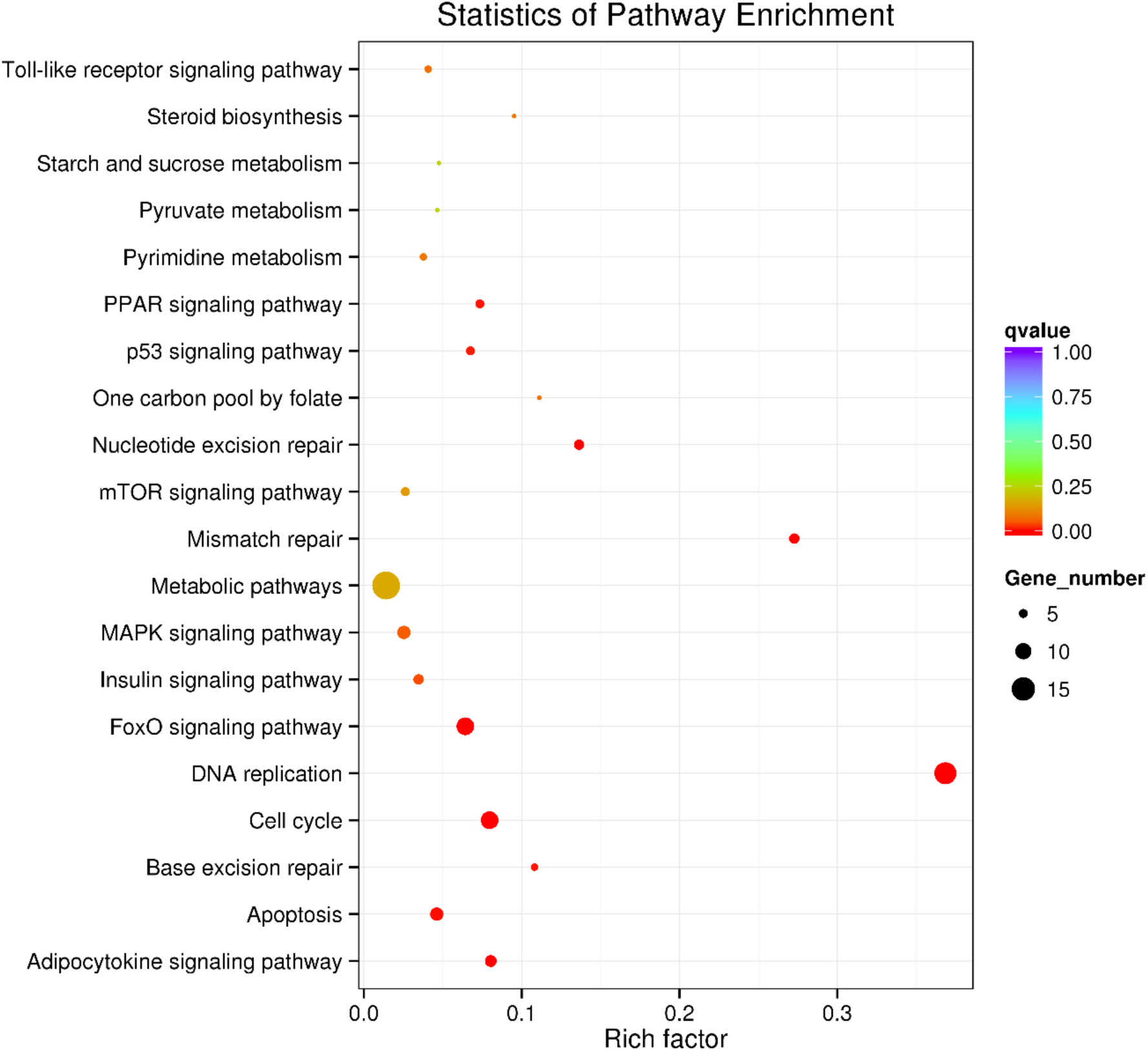
Overall KEGG pathway enrichment statistics for UH vs Veh.

## EXPERIMENTAL DESIGN, MATERIALS, AND METHODS

### Zebrafish husbandry and embryo collection

Adult wildtype AB strain zebrafish were maintained in Tecniplast ZebTec Active Blue^TM^ semi-recirculating aquatic habitat systems, located in a temperature-controlled room (26 °C ± 1) with a 14/10-hour light-dark cycle. System water conditions were continuously monitored and maintaining temperature, pH, and conductivity (28.5 °C ± 0.5, pH 7.5 ± 0.25, and 700 µSiemens ± 100, respectively). Fish were fed twice daily with Skretting Gemma® 75, 150, or 500, depending on size. Embryos were obtained by spawning adult zebrafish (≥120 dpf) in breeding tanks with perforated inserts that allow embryos to pass through but not adults. Embryos were collected from these breeder tanks and rinsed with fresh 0.3x Danieau’s solution (DanS), then moved to 15 cm petri dishes and maintained in an incubator set to 28.5 °C with an internal 14/10-hour light/dark cycle.

### Treatment solution preparation

Uranium in the form of uranyl nitrate (UN; (UO_2_(NO_3_)_2_) 6·H_2_O; CAS# 13520-83-7, Electron Microscopy Sciences, CAT# 22600), a soluble DU species, was used to prepare a 3 mg/mL stock solution in milliQ-water and diluted with DanS to produce treatment solutions of 30 and 300 µg/L UN or 18.1 and 181 ppb U atoms in 0.3x DanS, respectively. Sodium (meta) arsenite (NaAsO₂) was obtained from Sigma Aldrich (CAT# S7400) and dissolved in milliQ-water to produce a 1 mM stock solution. Next, a working solution was prepared by diluting the stock solution with milliQ-water and 30x DanS to a final concentration of 1 µM NaAsO₂, in 0.3x DanS.

### Waterborne larval exposures

At 4 hours post fertilization (hpf), larvae were moved into separate petri dishes containing treatment solutions of 30 or 300 µg/L UN, 1 or 10 µM NaAsO₂, or 1 µM NaAsO₂ and 30 µg/L UN in combination. Larva were maintained in these dishes with twice daily media changes until 5 dpf.

### Sample preparation and RNA extraction

Total RNA was extracted from pooled zebrafish larvae (n = 10 per biological replicate) using TRIzol reagent (CAT# 15596018; Thermo Fisher Scientific) and Direct-zol RNA MiniPrep Kit (CAT# R2050; Zymo Research Corporation) according to manufacturers’ protocols. RNA concentration and purity were assessed using a Nanodrop Lite (Thermo Fisher Scientific) spectrophotometer and RNA was qualitatively evaluated by absorbance ratios: A_260/280_ ratio between 1.8 and 2.2, A_260/230_ ≥ 1.8, and by visualization on a 1% agarose gel. Qualified RNA samples were shipped on dry ice to Novogene Co., Ltd. (Beijing, China) for library construction, sequencing, and basic differential gene expression analyses.

### RNA quality control and library preparation

RNA integrity was assessed using an Agilent 2100 Bioanalyzer. Samples exceeded the quality criteria of RNA Integrity Number (RIN) ≥ 5.8, smooth baseline, and total RNA yield ≥400 ng (**Supplemental Table 1**). Polyadenylated mRNA was enriched from total RNA using oligo(dT) magnetic beads and fragmented using a proprietary fragmentation buffer. First-strand cDNA synthesis was accomplished using random hexamer primers and reverse transcriptase, followed by second-strand synthesis using a proprietary Illumina second-stand synthesis buffer with dNTPs, RNase H, and *E. coli* DNA polymerase I to produce the second strand via nick-translation. The resulting cDNA underwent end repair, A-tailing, ligation with Illumina-compatible sequencing adapters (RNA adapter sequences are from TruSeq^TM^ RNA and DNA sample prep kits; RNA 5’ adapter, part #15013205 5’-AATGATACGGCGACCACCGAGATCTACACTCTTTCCCTACACGACGCTCTTCCGATCT-3’; RNA 3’ adapter part #15013207 5’-GATCGGAAGAGCACACGTCTGAACTCCAGTCAC-(6 nucleotide index)-ATCTCGTATGCCGTCTTCTGCTTG-3’), size selection, and PCR amplification. Libraries were then validated on an Agilent Bioanalyzer, quantified using a Qubit 2.0 fluorometer (Life Technologies), and further assessed by qPCR to ensure a minimum library activity of >2 nM.

### RNA sequencing

Qualified libraries were sequenced on an Illumina NovaSeq 6000 platform using 150 bp paired-end reads, resulting in an average of 58.9 million read pairs. Raw reads were filtered to remove adapter sequences, reads with >10% unknown bases (N), or >50% low-quality bases (Q-score < 20). Mapped reads corresponded to approximately 75% exonic, 20% intergenic, and 5% intronic DNA, +/- 3%. Clean reads were used for downstream analysis.

### Alignment, differential gene expression, and pathway enrichment analyses

Clean filtered reads were aligned to the *Danio rerio* reference genome GRCz11 using HISAT2 (v2.1.0) with default parameters [6]. Gene-level quantification was performed using HTSeq (v0.6.1) in union mode, and expression levels were normalized and reported as fragments per kilobase of transcript per million mapped reads (FPKM) [7]. The raw reads and FPKM statistics have been deposited in NCBI’s Gene Expression Omnibus [1] as GEO series accession number *GSE319292* (https://www.ncbi.nlm.nih.gov/geo/query/acc.cgi?acc=GSE319292) [2]. Differential expression analysis was performed using DESeq2 (v1.10.1) for comparisons with biological replicates, p-values were estimated using a negative binomial distribution, a significance threshold of adjusted p-value < 0.05 was applied, and false discovery rate was controlled using the Benjamini-Hochberg procedure [8]. Gene Ontology (GO) enrichment analysis was conducted using GOseq [9] and topGO [10], while KEGG pathway enrichment was performed using KOBAS (v3.0; [11]) against the KEGG database [12]. Plots were created using ggplot2 (v3.1.0+) in R Studio.

## LIMITATIONS

The approach using pooled larvae and total larval RNA sampling has several limitations. Heterogeneous transcript enrichment between individual larvae is unresolvable. Sequencing RNA from whole larvae may dilute significant cell-type specific responses, in some cases making them undetectable. Single-cell RNA sequencing would be ideal for overcoming those limitations. The current GRCz12tü reference zebrafish (Tübingen strain) genome assembly is available with increased annotated mRNAs, lncRNAs, and miscellaneous RNAs. However, our study was conducted in AB strain zebrafish with sequences mapped to the GRCz11 assembly, created from the transcriptomes of Tübingen crossed with AB fish [13].

## ETHICS STATEMENT

All zebrafish experiments were approved prior to execution by Northern Arizona University’s Institutional Animal Care and Use Committee (IACUC), protocol number 17-011. The methods and approaches followed animal usage guidelines as described in the NIH Guide for the Care and Use of Laboratory Animals [14]. No experiments involved human subjects or data collected from social media.

## CRediT AUTHOR STATEMENT

**Phillip Kalaniopio**: formal analysis, investigation, data curation, writing – original draft, review & editing, visualization. **Ronald Allen**: formal analysis, investigation, data curation. **Matthew Salanga**: conceptualization, methodology, investigation, formal analysis, resources, writing – review and editing, supervision, project administration, funding acquisition.

## ACKNOWLEDGEMENTS

This work was supported by NAU’s Technology Research Initiative Fund (TRIF; MCS), a pilot project award from the Southwest Health Equity Research Collaborative (SHERC; NIH 1U54MD012388-01, MCS) and support from the National Institutes of Environmental Health Sciences (NIEHS, 1R15ES032923-01/S1, MCS/PHK). Student support came from the Native American Cancer Partnership (NACP, NIH 1U54CA143925, PHK), NAU Louis Stokes Alliance for Minority Participation (LSAMP, NSF HRD1712523, PK), and the Research Initiative for Science Enhancement (RISE; NIGMS 1R25GM127199-01, RSA).

## DECLERATION OF COMPETING INTERESTS

The authors declare that they have no known competing financial interests or personal relationships that could have appeared to influence the work reported in this article.

## FIGURE LEGENDS

**Figure 1.** Volcano scatterplots of differentially expressed genes per condition versus vehicle control (1a-e), thresholds (dotted lines) set as p_adj_ > 0.05. Co-expression Venn diagram of all differentially expressed genes in and between all treatment groups (compared to vehicle control) (1f).

**Figure 2.** Overall KEGG pathway enrichment statistics for each condition versus vehicle control, with the top 20 most enriched pathways (or all pathways, if less than 20 are detected as enriched) on the y-axis and Rich factor on the x-axis. Q-value is indicated by the rainbow color scale of points, while size of each point is dictated by contributing gene number (2a-e).

