## Supplemental Table 1 for "Transcriptomic data of larval zebrafish exposed to continuous sub- and supra-MCL sodium arsenite and uranyl nitrate"

**Supplemental Table 1**. Sample RNA quality control provided by Novogene Inc., performed on an Agilent 2100 Bioanalyzer.

| Sample Name | Concentration (ng/μL) | Vol (μL) | Mass (μg) | RIN | Conclusion |
|---|---|---|---|---|---|
| DrVeh_1 | 175.33 | 23 | 4.033 | 10 | Pass |
| DrVeh_2 | 182.85 | 22 | 4.023 | 10 | Pass |
| DrVeh_3 | 181.62 | 24 | 4.359 | 9.5 | Pass |
| DrVeh_4 | 111.30 | 23 | 2.560 | 10 | Pass |
| DrAsL_1 | 230.31 | 23 | 5.297 | 9.9 | Pass |
| DrAsL_2 | 213.12 | 22 | 4.689 | 9.8 | Pass |
| DrAsL_3 | 122.66 | 23 | 2.821 | 10 | Pass |
| DrAsL_4 | 205.47 | 23 | 4.726 | 10 | Pass |
| DrAsH_1 | 201.38 | 28 | 5.639 | 9.9 | Pass |
| DrAsH_2 | 215.98 | 17.5 | 3.780 | 10 | Pass |
| DrAsH_3 | 207.57 | 20.3 | 4.214 | 9.9 | Pass |
| DrAsH_4 | 271.06 | 23 | 6.234 | 10 | Pass |
| DrAsU_1 | 209.18 | 19.5 | 4.079 | 9.9 | Pass |
| DrAsU_2 | 366.44 | 23 | 8.428 | 10 | Pass |
| DrAsU_3 | 259.88 | 19.6 | 5.094 | 10 | Pass |
| DrAsU_4 | 376.13 | 19.3 | 7.259 | 9.9 | Pass |
| DrUL_1 | 247.20 | 22 | 5.438 | 9.9 | Pass |
| DrUL_2 | 247.68 | 22 | 5.449 | 9.9 | Pass |
| DrUL_3 | 200.91 | 22 | 4.420 | 10 | Pass |
| DrUL_4 | 319.31 | 19.3 | 6.163 | 9.9 | Pass |
| DrUH_1 | 265.05 | 20.6 | 5.460 | 10 | Pass |
| DrUH_2 | 190.99 | 23 | 4.393 | 10 | Pass |
| DrUH_3 | 315.61 | 20.7 | 6.533 | 10 | Pass |
| DrUH_4 | 141.07 | 19.7 | 2.779 | 10 | Pass |
